# The United States Swine Pathogen Database: integrating veterinary diagnostic laboratory sequence data to monitor emerging pathogens of swine

**DOI:** 10.1101/2021.04.16.439882

**Authors:** Tavis K. Anderson, Blake Inderski, Diego G. Diel, Benjamin M. Hause, Elizabeth G. Porter, Travis Clement, Eric A. Nelson, Jianfa Bai, Jane Christopher-Hennings, Phillip C. Gauger, Jianqiang Zhang, Karen M. Harmon, Rodger Main, Kelly M. Lager, Kay S. Faaberg

## Abstract

Veterinary diagnostic laboratories annually derive thousands of nucleotide sequences from clinical samples of swine pathogens such as porcine reproductive and respiratory syndrome virus (PRRSV), Senecavirus A, and swine enteric coronaviruses. In addition, next generation sequencing has resulted in the rapid production of full-length genomes. Presently, sequence data are released to diagnostic clients for the purposes of informing control measures, but are not publicly available as data may be associated with sensitive information. However, public sequence data can be used to objectively design field-relevant vaccines; determine when and how pathogens are spreading across the landscape; identify virus transmission hotspots; and are a critical component in genomic surveillance for pandemic preparedness. We have developed a centralized sequence database that integrates a selected set of previously private clinical data, using PRRSV data as an exemplar, alongside publicly available genomic information. We implemented the Tripal toolkit, using the open source Drupal content management system and the Chado database schema. Tripal consists of a collection of Drupal modules that are used to manage, visualize, and disseminate biological data stored within Chado. Hosting is provided by Amazon Web Services (AWS) EC2 cloud instance with resource scaling. New sequences sourced from diagnostic labs contain at a minimum four data items: genomic information; date of collection; collection location (state or province level); and a unique identifier. Users can download annotated genomic sequences from the database using a customized search interface that incorporates data mined from published literature; search for similar sequences using BLAST-based tools; and explore annotated reference genomes. Additionally, because the bulk of data presently are PRRSV sequences, custom curation and annotation pipelines have determined PRRSV genotype (Type 1 or 2), the location of open reading frames and nonstructural proteins, generated amino acid sequences, the occurrence of putative frame shifts, and restriction fragment length polymorphism (RFLP) classification of GP5 genes. Genomic data from seven major swine pathogens have been curated and annotated. The resource provides researchers timely access to sequences discovered by veterinary diagnosticians, allowing for epidemiological and comparative virology studies. The result will be a better understanding on the emergence of novel swine viruses in the United States (US), and how these novel strains are disseminated in the US and abroad.

**Database URL:** https://swinepathogendb.org

## INTRODUCTION

The diversity of endemic viruses that circulate in swine continues to increase. Many of these viruses cause disease that adversely affect health and wellbeing through morbidity and mortality, and impact production through increased costs associated with vaccination, treatment, and increased biosecurity programs (1-3). In addition, novel viruses appear to periodically emerge in swine populations (e.g., Senecavirus A (SVA), porcine epidemic diarrhea virus (PEDV)) which can lead to the establishment and persistence of antigenically distinct viruses that lack appropriate vaccines due to limited information from which to derive rational formulations (4,5). Given that the genetic makeup of viruses continually changes, monitoring the patterns of evolution of viruses in swine is needed to identify possible emerging threats and to help control endemic viruses.

Veterinary diagnostic laboratories in the United States sequence thousands of clinical samples or isolates of porcine reproductive and respiratory syndrome virus (PRRSV), SVA, PEDV and other coronaviruses annually (4,6-17). The advent and broad use of next generation sequencing platforms has also resulted in the rapid production of full-length genomes. Unfortunately, these pathogen sequences are rarely publicly available, as the samples may be associated with sensitive information that veterinarians and pork producers may not want to disclose, or the process of annotating, validating, and sharing the genomic sequence information is burdensome. Currently, public genomic information is housed in NCBI GenBank (18), a comprehensive sequence database. Information from GenBank can be difficult to navigate for the purpose of identifying relevant sequence information and submitted data receives cursory curation and data may not be accurate. Ideally each nucleotide sequence would be annotated with start and stop sites for translation, the type of virus, potential open reading frames, along with the inclusion of data useful for genomic epidemiology studies, e.g., age of pig, collection location, collection date. Thus, swine disease researchers, outside of diagnostic laboratories, have limited means to identify novel isolates, where pathogens emerged, re-emergence of an identical but previously detected pathogen, and other knowledge that may be applied to improve animal health.

To address this problem for PRRSV, a producer funded initiative was implemented and maintained from 2005-2008 (Porcine reproductive and respiratory syndrome virus database - prrsvdb). The prrsvdb archived over 13,000 PRRSV orf5 sequences from both Type 1 (European) and Type 2 (North American) isolates from predominantly regional veterinary diagnostic labs and deposited over 8,200 unique sequence submissions to GenBank. The sequences generated and shared in this three-year period represent 25% of all available PRRSV data, included many critical PRRSV index strains derived from early 1990 field isolates. These data have been heavily used by molecular epidemiologists and other researchers worldwide and are still being accessed (19-22).

We developed the United States Swine Pathogen Database (US-SPD) to provide a mechanism for incorporating veterinary diagnostic data with sequence data for swine pathogens. The database was designed for the exploration of carefully curated genetic sequences to allow researchers and stakeholders the ability to determine how genetic diversity of swine pathogens is changing spatially and temporally. Sequences in the database are derived from public data in NCBI GenBank and clinical samples submitted to the Iowa State University Veterinary Diagnostic Laboratory (ISU VDL), the Kansas State University Veterinary Diagnostic Laboratory (KSU VDL), and the South Dakota Animal Disease Research & Diagnostic Laboratory at South Dakota State University (SD ADRDL). The core function of this database is to collect, store, view, annotate, and query genomic data for major swine pathogens, including PRRSV, SVA, PEDV, porcine deltacoronavirus (PDCoV), foot-and-mouth disease virus (FMDV), African swine fever virus (ASFV), and classical swine fever virus (CSFV). The apparent utility of genomic data in epidemiological analyses and control strategies (e.g., (23,24)) has facilitated a proliferation of genomic databases. Projects such as the Influenza Research Database (13) and ISU FLUture (14) implement curation and analytical tools for single pathogens.

The comprehensive NCBI viral genome resource provides stringent reference sequence annotation information. Some databases ingest and visualize public data, such as the ASFVdb (15). Our approach complements these by including customized curation pipelines for other swine pathogens, additional curation by subject-area experts (25), and by providing diagnostic laboratory data alongside public data. We additionally introduce a pathogen agnostic analysis pipeline, the swine pathogen analysis resource (SPAR), that accurately annotates sequence data with genomic features, and exports files that may be ingested into the US-SPD or NCBI GenBank.

## MATERIALS AND METHODS

### Database construction and implementation

The US-SPD operates as a web-based, curated, stable, relational database as part of infrastructure developed by the United States Department of Agriculture, Agricultural Research Service (USDA-ARS SCINet). The web interface of the US-SPD was developed using Tripal v3 (26-28) that builds upon the open source Drupal content management system and the Chado database schema (29). Tripal extension modules that incorporate NCBI BLAST sequence similarity search (27) and JBrowse genome browser (21,22) were implemented, providing useful tools for genetic data analysis and visualization. We modified the Tripal BLAST UI module to classify and visualize RFLP patterns of PRRSV orf5 genes. Additionally, we developed a custom Drupal module for the search interface that incorporates the ability to search genomic information derived from publications accessible via NCBI PubMed (30) or through metadata provided by sequence submitters in NCBI GenBank: these workflows and the search interface (e.g., search by gene name, date of collection, or sequence submitter) are in query pipelines in SQL. The database web server is hosted on an Amazon EC2 instance, currently a Linux server (Ubuntu 16.04 LTS), with Apache v2.4.18, PostgreSQL v9.5.23, PHP v7.0.33, and Drush v5.10. The US-SPD is updated monthly, and the curation pipeline is scheduled to run automatically when new data are acquired from diagnostic labs or public databases.

### Data collection and processing

All sequence data within the US-SPD has been curated following a genome annotation pipeline (available at https://github.com/us-spd) capable of identifying any kind of continuous genomic feature that can be translated into an amino acid sequence (Figure 1). In brief, the process can be broken down into two sections: pre-processing and query curation. Pre-processing consists of designing reference files necessary for annotating query sequences. A python script builds a reference scaffold multiple sequence alignment (MSA), and companion profile hidden Markov model (HMM) and general feature format (GFF3) templates. The MSA is then used to derive a series of JSON files including regular expression patterns that represent spaced fragments of genomic features. When the requisite files have been generated, query sequences may be submitted for curation. First, regular expressions from the JSON reference are used to determine position and reading frame of genomic features. If the regular expressions are unable to annotate submitted sequences, profile HMM alignment is performed to supplement pattern matching in order to reliably find feature terminal ends and/or frameshift locations (Figure 2). The curation pipeline can also use BLAST to identify query species or, in the case of PRRSV, genotype. Deployment of the annotation pipeline occurs in conjunction with parsing scripts that derive genomic information and metadata such as: sample source; collection date; collection location; NCBI PubMed ID; Authors; and strain name (Figure 2).

**Figure 1.**
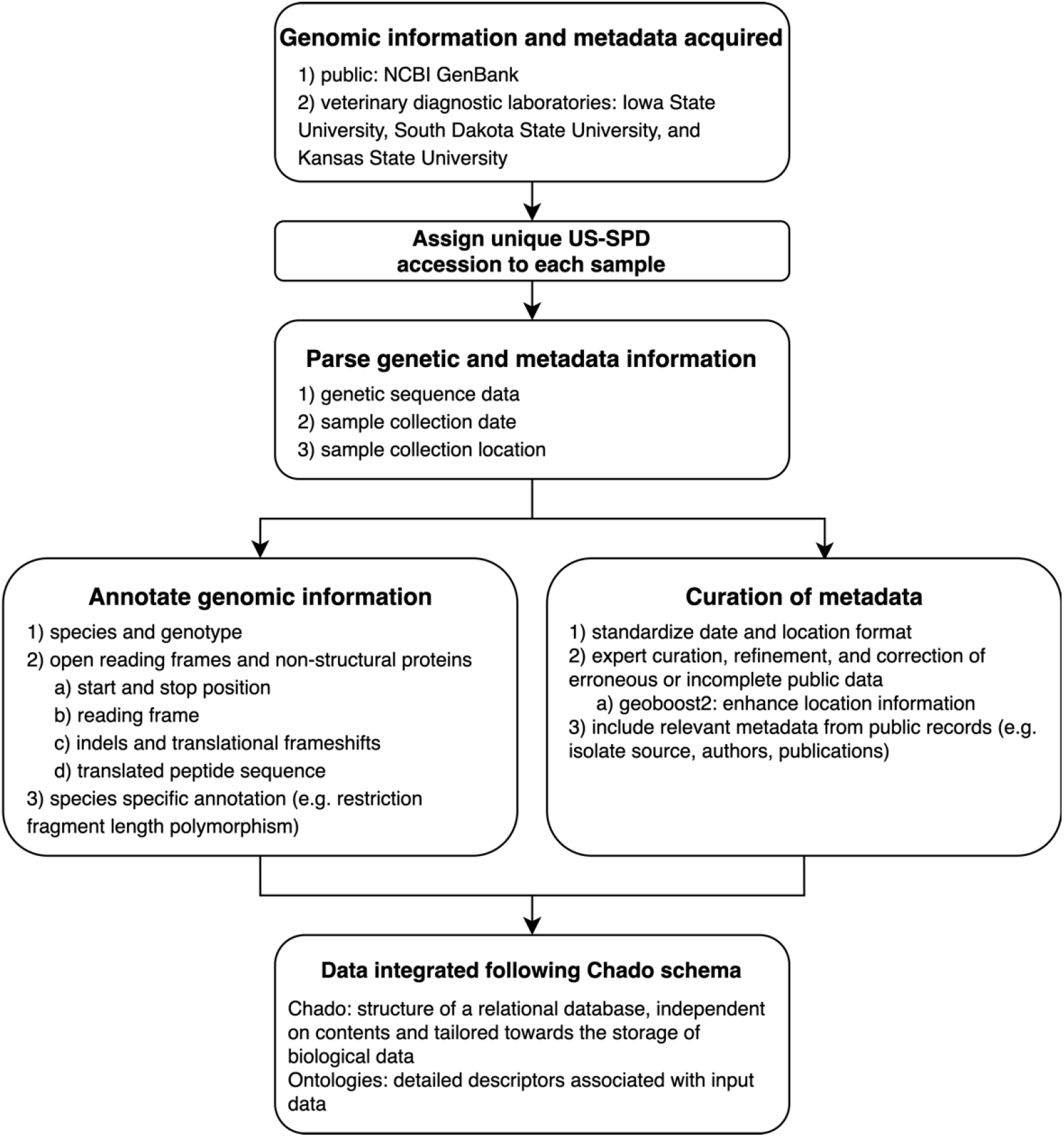
Conceptual model describing the automated pipeline implemented in the US Swine Pathogen Database that takes raw sequence data to fully anonymized and annotated virus sequence record in the relational database.

**Figure 2.**
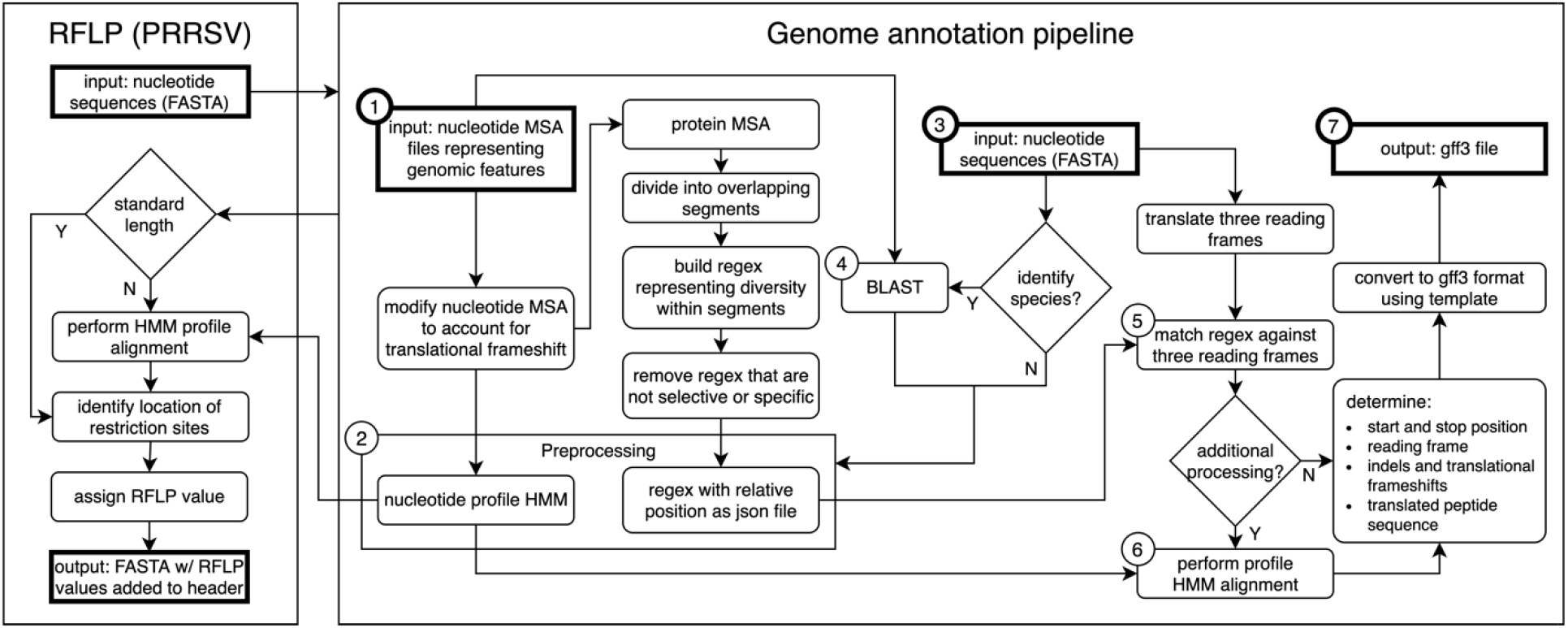
Genome annotation for the United States Swine Pathogen Database. Genome annotation begins with preprocessing, which requires nucleotide multiple sequence alignment files (MSA) representing genomic features as input (➀). The products of preprocessing (➁) are a nucleotide profile hidden Markov model (HMM) and a structured file containing regular expression patterns representative of diversity within sections of translated input MSAs. Following preprocessing, query nucleotide sequences in FASTA format (➂) may be supplied to the annotation pipeline. If species identification is necessary (e.g., differentiating type 1 and 2 PRRSV), BLAST is performed (➃) using preprocessing input files (➀) as reference. Once the query sequence species is known, relevant preprocessing files (➁) are selected. Regular expressions are matched against three reading frames (➄) of query sequence (➂) to determine the location of genomic features. If more processing is necessary due to frame changes or uncertainty in the start or stop position, a profile HMM alignment is performed (➅). These steps produce genome annotation and additional information with high confidence. The output produced by the pipeline is general feature format (GFF3) file, a standard nucleic acid or protein feature file format (➆). The annotation pipeline is available at https://github.com/us-spd/

At the core of the annotation pipeline are template nucleotide MSAs that are representative of the genetic data of each swine pathogen. The MSAs are used to generate amino acid (AA) regular expressions and nucleotide profile HMM templates and through this process are robust and prevent curation failure by capturing as much genetic variation as possible. A consequence is that the AA regular expressions within the pipeline may become very obtuse in areas where homology is low. For example, in PRRSV it was not uncommon for 10+ unique residues to be observed at any given position, which produces a pattern that may not match selectively. To improve selectivity, low occurrence residues may be removed from the automated patterns until the expression is passably unique.

The outcome is highly curated genomic information that allows users to search and download data for additional user applicable analyses that would not have been possible without the annotation process (e.g., fine-scale analysis of the spatial dissemination of viruses within the United States). For data provided by the veterinary diagnostic laboratories, the annotation tool produces a GFF3 formatted file that can be uploaded to NCBI GenBank for public dissemination of the data with appropriate recognition given to the individual laboratory.

### Customized search interface for genomic epidemiology and comparative virology

Genomic epidemiology and comparative virology rely on four core data items: genomic information; the date of collection; the US state or country the collection was made; and a unique identifier. The US-SPD system standardizes data from three major veterinary diagnostic labs for swine and NCBI GenBank. Newly ingested data from the diagnostic labs curated data by the US-SPD has this information, and the ability to exclude sequences that do not meet these criteria (Figure 1). In the interests of diagnostic laboratory confidentiality issues, diagnostic laboratory data that has been submitted to the US-SPD has an anonymized unique ID, i.e., only the submitting laboratory retains the ability to track a sample and its genomic information and metadata to a producer or veterinarian. The US-SPD has also ingested and curated publicly available sequence data; in these cases, custom parsers have extracted available metadata, but these records may not have information beyond sequence.

The US-SPD custom search interface allows users to query and retrieve data in standard formats, e.g., FASTA sequence file, comma separated text files with associated metadata. The search interface is split by pathogen, and queries may be based on: virus; gene; date of collection; state of collection; isolation source; sequence length, completeness, with duplicate records identified; Author or submitting laboratory; PubMed ID; and isolate strain name. This returned information is standardized and with additional analyses can be applied for vaccine design; determining when and how fast pathogens are spreading across the landscape; genomic structure determination; and translational and post-translational protein analyses. Importantly, by integrating large-scale genomic information, representative virus genes or genomes may be selected, potentially sourced from regional diagnostic labs, and used in comparative approaches to determine phenotypic consequences of genetic diversity, e.g., whether specific nucleotide and/or amino acid changes affect phenotype (31). Whole genome strain data is searchable using similar strategies. For viruses such as PRRSV, this provides the ability to study the role of recombination in evolutionary dynamics, and identify vaccine targets outside of the structural genes, e.g., nonstructural protein 2 (32,33).

### Identifying homologous sequence data, PRRSV orf5 classification, and gene visualization

The US-SPD includes an implementation of the Tripal BLAST UI extension module (https://github.com/tripal/tripal_blast/). This module is an intuitive implementation of the NCBI Blast+ tools (34), and relies upon the Tripal interface with Drupal integration for easy deployment and management. As applied in the US-SPD, the tool provides access to nucleotide- and protein-based BLAST functionality (e.g., blastn, blastx, blastp, tblastn) and is integrated to facilitate rapid queries and job submission for all pathogens with a jobs daemon. The US-SPD currently allows queries against all data in the database, or specific subsets of that data, i.e., a user is able to query the aggregated database, or a specific pathogen, or annotated sequences. The utility of querying annotated sequences is most apparent with PRRSV, whereby the similarity of a query orf5 gene may be determined against an established orf5 genetic nomenclature that assigns lineage information (35). The module produces a graphical exploration of genetic similarity, including the reference sequence location where the query gene is most similar. These data can be used to locate similar genes within the database for further comparative analyses or to quickly identify novel viruses that have not been previously detected in US swine, e.g., (36).

Custom analysis scripts have been deployed through modification of the Tripal BLAST UI extension module. Specifically, a script rapidly classifies PRRSV virus 2 strains by restriction fragment length polymorphism (RFLP) patterns from user submitted query sequences and outputs corresponding RFLP assignment. RFLP values have been extensively used as a means to quantify the diversity of orf5 genes (e.g., (21,22)). The approach uses cut site locations created by three restriction enzymes (MluI, HincII, and SacII) to classify strains based on the cut site of each restriction enzyme. Although the application of this tool has limitations as diversity in orf5 may not reflect diversity across the genome or the phenotype of a virus, it has utility in assessing, categorizing, and linking contemporary genetic signatures to archived sequence records. The implemented tool takes user-submitted orf5 nucleotide data, checks the length of the sequence and then aligns with HMMER3. Subsequently, the tool identifies restriction sites and classifies the viral sequence with the appropriate three-digit numeric code. RFLP patterns for PRRSV-2 orf5 sequences have been determined and stored within the database as a searchable field, allowing users to quickly screen their data and identify similar sequences for analyses.

To further facilitate comparative analyses of genomes, we have implemented the Triapl JBrowse extension module (https://github.com/tripal/tripal_jbrowse). The module embeds JBrowse (22), an interactive, client-side genome browser, into a Drupal webpage and provides a simple interface for managing and creating JBrowse instances. We have deployed seven JBrowse instances covering ASFV, CSFV, FMDV, PDCoV, PEDV, PRRSV, and SVA with annotations of coding sequence, genes, mature peptides, and untranslated regions of reference sequences from the NCBI RefSeq database (http://www.ncbi.nlm.nih.gov/genome). This feature remains in active development and may be refined as more information is derived during sequence annotation, incorporated through subject-area expert input, and these metadata may be captured and shared within the JBrowse interface. As this tool is integrated within the Tripal toolkit, future versions of the US-SPD have the potential to integrate user submitted information from each of the JBrowse genome instances into the database, or users can save and share their JBrowse annotations.

## UTILITY AND DISCUSSION

### Overview of the genomic data in the US-SPD

The US-SPD now includes 63,804 validated virus genomes, including 60,726 sourced from NCBI GenBank and an additional 3,078 from regional diagnostic labs. There are seven viruses currently in the database, each of which represents a major agricultural pathogen: ASFV, CSFV, FMDV, PDCoV, PEDV, PRRSV, and SVA. Each genome includes a link to the original GenBank record, and the information parsed from this record can be annotated onto the FASTA definition line or in a metadata download for additional analyses. Genes and/or strains that have been used in empirical studies have been incorporated for all viruses and are linked via the PubMed ID to the publication. In total, there are 190,828 validated viral and genome segments within the US-SPD database. The increase of data availability reflects the detection of novel viruses (23,24,37-43). Unfortunately, while some of the virus strains in the US-SPD are well studied in the laboratory, many other sequenced viruses and strains are not. The establishment of the US-SPD highlights the diversity of these viruses and provides an interface to study their phenotype.

### Cocirculation and turnover of multiple genetic clades of PRRSV

Porcine reproductive and respiratory syndrome virus (PRRSV) represents the single most significant threat to the health and economic productivity of swine in the United States today. Since its discovery in the late 1980s, PRRSV has caused devastating production losses in herds throughout the United States and the world (44-46). Despite significant advances in our understanding of PRRSV biology (47), the disease has remained very difficult to control. There are two reasons for the remarkable resistance of PRRSV to control through vaccination: firstly, PRRSV is immunomodulatory and immunoevasive; and secondly, PRRSV has extraordinary genetic and antigenic variability. PRRSV may suppress early events in the activation of host cellular immunity and misdirects or delays the production of neutralizing antibodies (47-49). Underlying this property is extraordinary genetic and antigenic variability, and a consequent ability to evolve rapidly due to the low fidelity of the viral RNA polymerase (50,51).

Using the US-SPD, we have generated a high-quality sequence dataset of PRRSV orf5 sequences that capture the extent of genetic diversity from 1990 to 2021. These data include all sequences from isolates in GenBank (n=29,325), alongside previously unpublished data derived from participating regional diagnostic labs (n=3,078). As of March 2021, these sequences were comprised of 32,403 worldwide samples with most viruses from the United States of America, but viruses from Canada, Mexico, China, Korea, Japan, Thailand, Austria, Denmark, Italy and Poland were also present. Using the RFLP-typing tool on the US-SPD, we quantified RFLP values for all complete sequences from 2000 to present as a means to quantify temporal changes in the diversity of orf5 genes. We document concurrent circulation of the 15 major RFLP patterns consistently across all years, i.e., patterns that were the most frequently detected in total across the time period, and an additional 138 “Other” patterns that represent ∼10% to ∼25% of detections each year representing 1991 total detections of 16403 total patterns (Figure 3). The predominance of certain RFLP patterns rapidly changes from year to year, e.g., 1-4-4 was ∼50% of detections in 2011-12 (declining to ∼10% in 2015), and was superseded by 1-7-4 type orf5 genes in 2015, but this RFLP pattern rapidly declined to ∼15% of detections by 2020. In the last two years, these data also revealed that no single RFLP pattern became predominant, with almost even detections of 1-7-4, 2-5-2, 1-8-4, 1-2-2, and “Other.”

**Figure 3.**
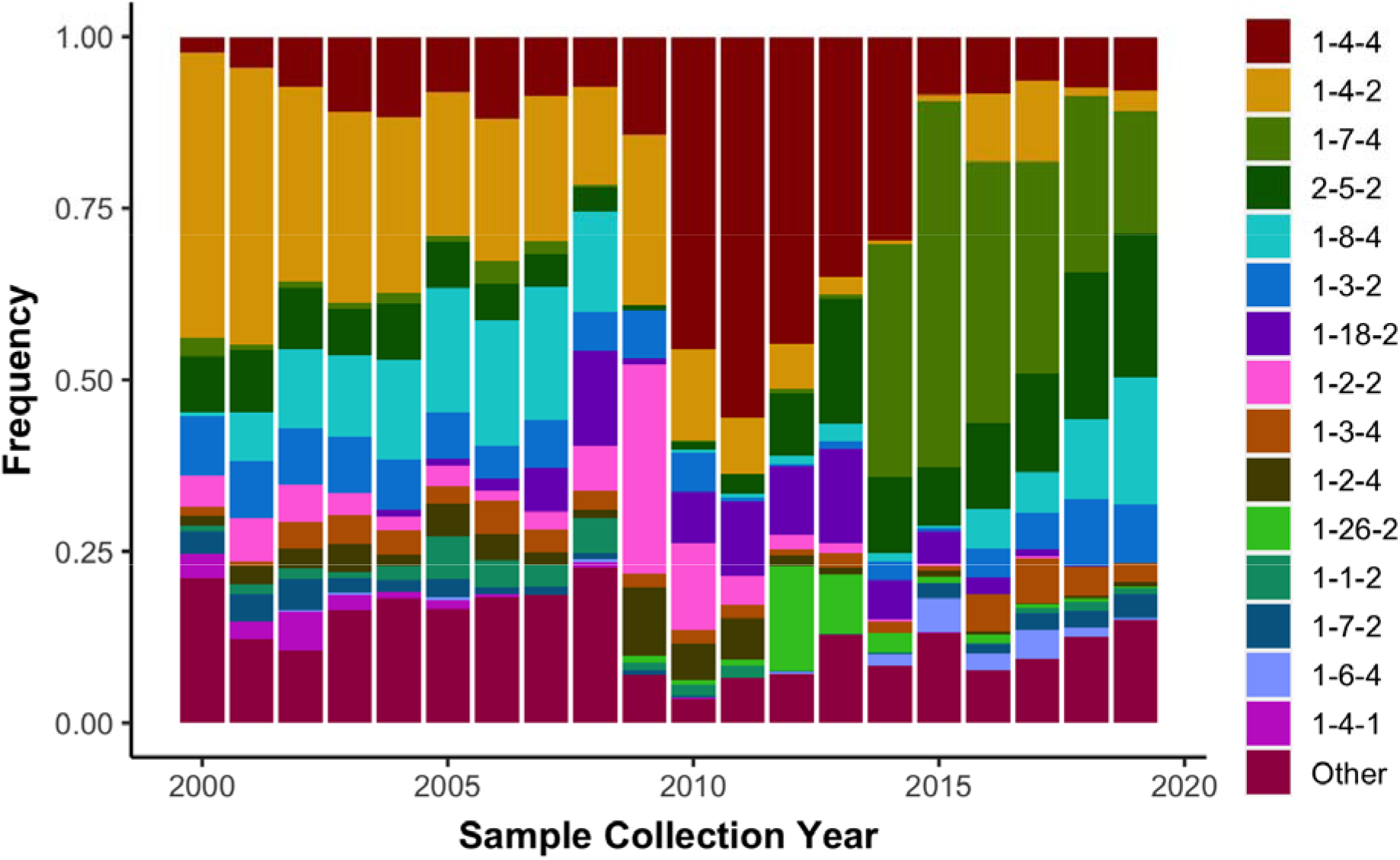
Observed frequency of the top 15 most commonly detected restriction fragment length polymorphism (RFLP) patterns in type 2 porcine reproductive and respiratory syndrome virus sampled in the United States from 2000 to present (n = 16,403). Less common RFLP patterns were grouped together and labelled as “Other”.

### Recombination and PRRSV evolution in the USA, 2014 to 2021

We demonstrate the utility of our search interface to conduct genomic analyses by conducting a phylogenetic network analysis of a field-relevant PRRSV dataset accounting for recombination. This is motivated by prior research (21) that demonstrated recombination in PRRSV resulted in differential evolutionary dynamics that were associated with unique patterns of pathogenesis and transmission.

Importantly, recombination in PRRSV has two biological outcomes: it increases genetic diversity; and it increases the likelihood of the emergence of a virus that is antigenically novel via a ‘sampling effect’, e.g. (59-62). In support of this hypothesis are data that demonstrate that following recombination, novel viruses with unique phenotypes have emerged in China, Europe, and the USA (21,58,63-67).

We apply a recent algorithm, RF-Net, that was built to infer virus networks accommodating for influenza A virus reassortment (53,54). This analytical method is derived from hybridization network analyses (52) and is similar to the inference of rooted phylogenetic trees but includes nodes that have more than one parent that can account for recombination. Functionally, the RF-Net algorithm synthesizes a collection of gene trees, and then embeds those gene trees into a network that displays each gene tree with the minimum number of recombination nodes (53,54).

For these analyses, we searched for all PRRSV complete strains (n=1,389) in the US-SPD. Completeness was determined by searching for the presence of all full open reading frames and non-structural proteins. We then restricted and identified strains that were collected in the USA from 2014 to present and downloaded the individual annotated genes (n=253). The genomes were separated into the constituent genes, and a single outgroup was included, VR-2332 (GB000137/DQ217415). To generate the required input trees for RF-Net, genes were aligned using default parameters in mafft (55), and maximum likelihood phylogenetic trees were inferred following automatic model selection using IQ-TREE (56,57). To infer the phylogenetic network, we applied the RF-Net algorithm and explored virus networks with different numbers of recombination events, *r*. Though there is an automatic stopping criterion when the improvement in the embedding cost is negligible, we chose to explore networks with *r* ranging from 0 to 20, and subsequently determined whether biologically plausible recombinant strains were detected (Figure 4).

**Figure 4.**
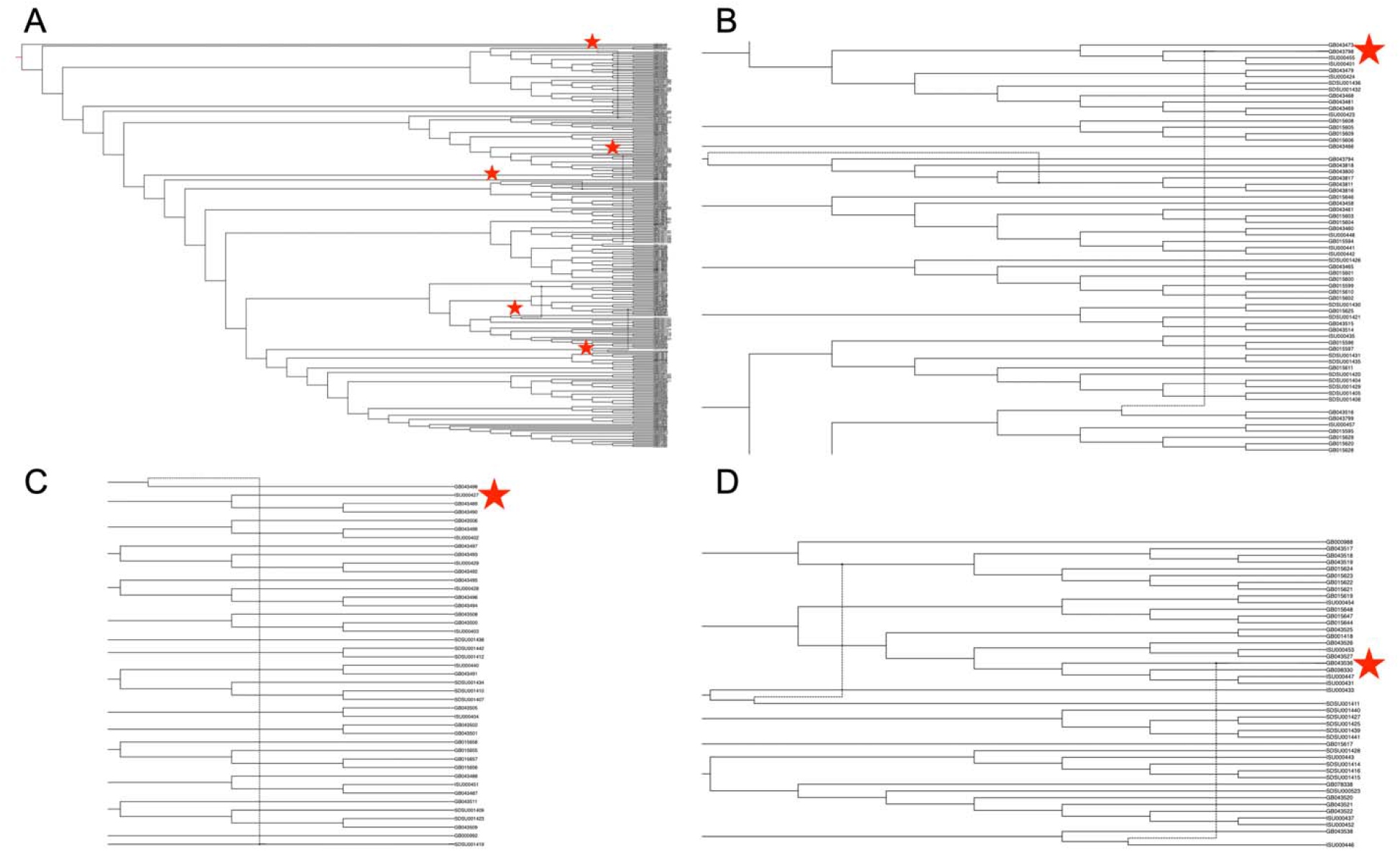
Phylogenetic network of porcine reproductive and respiratory syndrome virus collected in the Midwest of the United States from 2014 to present (A). Putative recombination events are indicated by vertical lines within the network, annotated by red stars. Panel (B) strain IA30788-R (GB043798), Panel (C) strain 23199-S4-L001 (GB043498), and Panel (D) strain 7705R-S1 (GB043536) are visualized separately to demonstrate evolutionary relationships and recombination nodes. Phylogenetic network with tip labels, recombination layers 1 through 20, and gene tree embeddings is provided at https://github.com/us-spd/.

We identified a network with a minimum of 3 recombination events as the best of those we explored, i.e., though embedding cost was minimized with *r* = 20 (data available at https://github.com/us-spd/), almost all of these recombinant events have not been characterized (but see (68,69)). Consequently, we visualize the network with *r* = 5 to demonstrate the 3 previously reported signals of recombination, alongside an additional 2 undescribed recombination events to highlight how unique strains may be identified (Figure 4). RF-Net identified strain IA30788-R (GB043798, Figure 4B) that has recently been described as a putative recombinant between a wildtype strain (IA76950-WT) and a commercial vaccine (70). In this case, our inference excluded the suggested parental strain (P129, GenBank: AF494042), but RF-Net was able to detect an evolutionary signal associated with recombination, i.e., RF-Net can use incomplete sequence data and reveal significant biological events. Similarly, additional recombination signals were detected in 23199-S4-L001 (GB043498, Figure 4C) and 7705R-S1 (GB043536, Figure 4D) strains that have also been reported as, or associated with, recombination (58). A further application of this inferred phylogenetic network is to generate a list of strain rankings based upon genetic diversity. Typically, PRRSV phenotypic studies focus on reference strains because of ease-of-access, publication history and evaluation of whether the variant represents a particular geographic location or biological property. Strains in networks such as these may be objectively ranked using algorithms such as the Fair Proportion Index or the Shapley Value (71-74), and strains that exhibit recombination signals, may have genetic features that require additional phenotypic characterization.

## Conclusions

The database and curation pipeline provides a recording system that will standardize data from the major swine veterinary diagnostic labs alongside public data deposited to NCBI GenBank. Further, the search interface allows the dissemination of these data in standard format (e.g., FASTA sequence file) based upon user queries.

This is a simple interface, but because the interface allows researchers to screen data rapidly it has the power to provide information for vaccine design, determining when and how fast pathogens are spreading across the landscape, identifying transmission hotspots at a coarse spatial scale, and generating datasets appropriate for comparative virology. All these analyses benefit from large-scale genomic information and these data have not been available or have been addressed using small – and perhaps inappropriate – datasets, e.g., (78). Additionally, through the standardized availability of date collection on all data, users may create time-scaled phylogenies allowing the time of emergence for novel viral isolates to be determined, and these data can be used to determine viral spread across the landscape using the state level geographical information provided by the diagnostic laboratories, e.g.,(58,76). Another benefit of these data is that genomic information covering thousands of sequences allows for the identification of critical amino acid substitutions associated with particular genetic clades of viruses. Understanding these critical substitutions can be used to inform vaccine updates and composition, e.g., (19,75).

### Future directions

As the rate of viral genome sequencing in swine virology increases, e.g., (77), databases are required to link those in the diagnostic community with researchers to improve animal health and economic productivity. This process should include careful curation, with standardized sequence annotation methods that ensure data quality to enhance our ability to make sound inference from the data, i.e., linking nucleotide/amino acid changes across disparate parts of the genome (epistasis), where insertions and deletions are evolving, where other key translation events are located, and linking genetic diversity to antigenic phenotype. Once annotated, large-scale genome sequence data needs to be available in ways that facilitate discovery. This requires automated metadata capture and data standardization, as well as interfaces that leverage the annotated metadata for scientific discovery. Many different approaches are possible (e.g., NCBI Virus Variation Resource (79)), but the US-SPD achieves this by providing search fields that are specific for the swine health community, and will allow for timely access to novel sequences discovered by regional veterinary diagnostic laboratories. While currently limited to seven swine pathogens, our intent is to expand the automated US-SPD data annotation pipeline to include more viruses, and also be flexible enough to account for the emergence of novel pathogens. As a public resource, the US-SPD serves a range of users in the animal health community (to date, registered users include regulatory agencies, industry, and academia) who conduct work in diagnostic labs developing assays, to those who conduct basic research in molecular virology to understand the biology, evolution, and transmission of viruses. Our mission is informed by their use, and by engaging our stakeholders and working together on shared goals we can provide the rigorous resources necessary to support a comprehensive swine pathogen database. Collectively, reducing the impact of viral pathogens in swine requires a fundamental knowledge of what viruses are circulating in the population and the US-SPD achieves this by providing a tool for investigators to develop rational and representative vaccines which will reduce viral burdens, decreasing the economic burden of viral disease and improving animal health through targeted interventions and surveillance.

## AVAILABILITY

The analytical code used in this research is available in the GitHub repository (https://github.com/us-spd/); the database implements the open source Tripal project which is available at the following website https://tripal.info/. Restrictions apply to the immediate availability of diagnostic laboratory data due to client confidentiality: these data are embargoed from 12 months from receipt from the diagnostic lab, and then become publicly available. Data may be available upon reasonable request and with permission of the contributing diagnostic laboratories.

## ACKNOWLEDGEMENTS

We gratefully acknowledge pork producers, swine veterinarians, and laboratories for participating in surveillance for RNA pathogens infecting swine and publicly sharing sequences in the United States Swine Pathogen Database and NCBI GenBank. The authors acknowledge the efforts in pathogen sequence generation and reporting by all technical personnel at Iowa State University Veterinary Diagnostic Laboratory, Kansas State University Veterinary Diagnostic Laboratory, and the South Dakota Animal Disease Research and Diagnostic Laboratory. We thank Alexey Markin for providing an pre-release version of RF-Net and discussing implementation and visualization of inferred virus networks.

